# Modulation in Alpha Band Activity Reflects Syntax Composition: An MEG Study of Minimal Syntactic Binding

**DOI:** 10.1101/2021.07.09.451797

**Authors:** Sophie M. Hardy, Ole Jensen, Linda Wheeldon, Ali Mazaheri, Katrien Segaert

## Abstract

Successful sentence comprehension requires the binding, or composition, of multiple words into larger structures to establish meaning. Using magnetoencephalography, we investigated the neural mechanisms involved in binding at the syntax level, in a task where contributions from semantics were minimized. Participants were auditorily presented with minimal sentences that required binding (pronoun and pseudo-verb with the corresponding morphological inflection; “*she grushes*”) and pseudo-verb wordlists that did not require binding (“*cugged grushes*”). Relative to no binding, we found that syntactic binding was associated with a modulation in alpha band (8-12 Hz) activity in left-lateralized language regions. First, we observed a significantly smaller increase in alpha power around the presentation of the target word (“*grushes*”) that required binding (-0.05s to 0.1s), which we suggest reflects an expectation of binding to occur. Second, during binding of the target word (0.15s to 0.25s), we observed significantly decreased alpha phase-locking between the left inferior frontal gyrus and the left middle/inferior temporal cortex, which we suggest reflects alpha-driven cortical disinhibition serving to strengthen communication within the syntax composition neural network. Together, our findings highlight the critical role of rapid spatial-temporal alpha band activity in controlling the allocation, transfer and coordination of the brain’s resources during syntax composition.

---

The expressive power of human language is largely derived from our ability to combine multiple words into larger syntactic structures with more complex meaning. Such binding, or compositional, processes occur during the comprehension of even the most basic two-word phrases (e.g., “*she walks*”). Characterizing the neural processes involved in composition – also referred to as *Unification* (Hagoort 2003) or *Merge* (Chomsky 1995) – has been a central topic of research for many years ((Hagoort 2019; Pylkkänen 2019; Matchin and Hickok 2020). In the present study we provide novel insight into the neural mechanisms involved in syntax composition using a task in which syntactic binding is dissociable from semantic composition. We use the term *syntactic binding* to specifically refer to the neural processes involved in the combining of individual words into larger structures. We employed a minimal phrase paradigm involving pseudo-words (i.e., following the phonotactic rules of a language, but not conveying semantic meaning).

Sentential compositional processes predominantly occur within a left-lateralized network of brain regions, including the inferior frontal gyrus (IFG) and angular gyrus (Friederici et al. 2000; Humphries et al. 2006; Pallier et al. 2011; Matchin et al. 2017). Within this network, modulations in theta, alpha and beta frequencies are thought to be crucial for higher-order linguistic functions (Bastiaansen et al. 2010; Meyer 2018; Prystauka and Lewis 2019). However, the precise neural mechanisms of syntax composition, relating to frequency modulations, remain elusive. This is because while some studies have found compositional processing to be associated with increased alpha and beta power (Meyer et al. 2013; Segaert et al. 2018), others have found sentence unification to be associated with a power *de*crease (Wang et al. 2012; Lam et al. 2016; Gastaldon et al. 2020). Moreover, it is unclear how functional connectivity (i.e., phase-locked) between the neural oscillations in different brain regions may contribute towards successful syntax composition. Current evidence suggests that functional connectivity between regions implicated in compositional processes, such as the left IFG, anterior temporal lobe (ATL) and posterior superior temporal gyrus, is beneficial for sentence comprehension (Schoffelen et al. 2017; Vassileiou et al. 2018; Lopopolo et al. 2021). However, the studies discussed thus far have typically used complex sentence structures – this means that other cognitive processes, such as working memory, are also involved in comprehending the sentence stimuli (Pylkkänen 2019), thereby making it difficult to functionally isolate the neural processes involved in sentence composition alone. In this study, therefore, we aim to characterize the neural mechanisms of syntax composition, both in terms of power modulations and phase-locked connectivity, at its most basic two-word level.

A short two-word sentence (e.g., “*red boat*”) is a traceable linguistic unit which can be used to decompose the brain networks implicated in the processing of lengthier sentences (Pylkkänen 2019). Research involving minimal sentences has identified a network of left-lateralized brain areas that underlie composition, most notably including the ATL (Bemis and Pylkkänen 2011, 2013; Pylkkänen et al. 2014; Westerlund et al. 2015; Zhang and Pylkkänen 2015; Ziegler and Pylkkänen 2016; Zaccarella et al. 2017; Schell et al. 2017; Blanco-Elorrieta et al. 2018; Zhang and Pylkkänen 2018; Fyshe et al. 2019; but cf. Kochari et al. 2021). Importantly, in order to more precisely identify the neural processes involved in syntactic binding that are dissociable from semantics, one further approach is to use pseudo-words within a minimal phrase paradigm. Using fMRI, Zaccarella and Friederici (2015) found increased hemodynamic responses in the anterior part of the left pars opercularis (part of the IFG) during comprehension of determiner-noun phrases involving pseudo-nouns (“*this flirk*”) compared to wordlists involving one pseudo-noun (“*apple flirk*”). Building on this, using EEG Segaert et al. (2018) found increased alpha and beta power (centralized over frontal-central electrodes) during comprehension of minimal phrases involving pseudo-verbs (“*she grushes*”) compared to wordlists of two pseudo-verbs (“*cugged grushes*”), which they interpreted as reflecting syntactic binding (see also Poulisse et al. 2020). We aim to build on these findings and use magnetoencephalography (MEG) to precisely characterize and localize the rapid temporal features and functional connectivity within a spatially distributed network of brain regions that support syntax composition independent of semantics.

We focused on frequency ranges up to 30 Hz as it is modulations in these frequencies that are considered critical for syntax composition (e.g., Bastiaansen et al. 2010; Segaert et al. 2018; Poulisse et al. 2020; Gastaldon et al. 2020); whereas semantic composition appears to involve higher gamma band activity (Hald et al. 2006; Bastiaansen and Hagoort 2015). In particular, we hypothesize to observe effects in the alpha (~8-12 Hz) frequency band given previous evidence that modulations in the alpha band reflect sentence compositional processing; however, the directionality of this effect is difficult to predict given that some research has found compositional processing to be associated with increased alpha (Meyer et al. 2013; Segaert et al. 2018), whereas others have found it to be associated with alpha decreases (Wang et al. 2012; Lam et al. 2016; Gastaldon et al. 2020). We may also reasonably expect to observe effects of syntax composition in the theta (~4-7 Hz) and beta (~15-30 Hz) frequency bands if they also contribute towards sentence comprehension (Meyer 2018; Prystauka and Lewis 2019). In line with previous work, we predominantly expect to observe oscillatory effects of syntactic binding immediately preceding and following the onset of the target word where binding occurs (i.e., “*grushes*”) (Bemis and Pylkkänen 2013; Segaert et al. 2018; Zhang and Pylkkänen 2018; Poulisse et al. 2020). We hypothesize that our observed oscillatory effects will be localized to left hemisphere language regions, including the LIFG (e.g., Hagoort 2003; Tyler and Marslen-Wilson 2008; Zaccarella and Friederici 2015; Zaccarella et al. 2017). However, the more interesting hypotheses relate to the functional connectivity between different language-relevant regions during syntactic binding. In particular, alpha band desynchronization has been found to predict successful sentence encoding (Magazzini et al. 2016; Vassileiou et al. 2018; Alavash et al. 2021), suggesting that alpha desynchronization serves to strengthen communication within a cortical network by enabling the increase transfer of information (Jensen and Mazaheri 2010; Klimesch 2012; Van Diepen et al. 2019). Our study is the first to closely examine functional connectivity among brain regions involved in minimal syntactic binding, but building on previous work, we may predict to observe less alpha-locking during syntactic binding compared to no binding.

## Materials and Methods

The methods and planned analyses of this study were pre-registered on the Open Science Framework prior to data collection (https://osf.io/ntszu).

### Participants

We recruited 25 healthy participants: all were right-handed and native monolingual British-English speakers. One participant was excluded due to excessive movement artefacts during the MEG test session (> 50% trials removed), meaning that a sample of 24 participants was used in the time-frequency analyses (13 female / 11 male, *M* = 24.2yrs, *SD* = 4.1yrs). Anatomical T1 brain scans were acquired for 21 of the participants (two participants did not attend the MRI session, and another did not complete the MRI session due to unexpected discomfort). Further technical issues with MRI-MEG co-registration with two participants meant that a sample of 19 participants was used for the source localization and connectivity analyses (10 females / 9 males, *M* = 24.1yrs, *SD* = 4.2yrs). The study was approved by the University of Birmingham Ethical Review Committee. All participants provided written informed consent and were compensated monetarily.

### Experimental Design and Stimuli

We employed a simple design of two experimental conditions: the sentence condition consisting of a minimal two-word phrase (pronoun plus pseudo-verb) for which it was highly likely that syntactic binding may plausibly occur (e.g., “*she grushes*”); and the wordlist condition consisting of two pseudo-verbs for which syntactic binding was highly unlikely to occur (e.g., “*cugged grushes*”). Syntactic binding occurred in the sentence condition (but not the wordlist condition) because the correct morphological inflection (i.e., -es) cued binding with the corresponding pronoun, in a way that is similar to how an intransitive verb phrase may be interpreted (e.g., “*she grushes*” is akin to “*she walks*”). Varying the first word between the sentence and wordlists conditions (while ensuring that the second word was matched across the two conditions) enabled us to experimentally manipulate the binding context of the second word (syntactic binding vs. no binding). Our analyses therefore specifically focus on the neural signature surrounding the presentation of the second word only as this is the time period of interest where we expect binding to occur, matching the approach taken in other minimal binding studies (Bemis and Pylkkänen 2011, 2013; Zaccarella and Friederici 2015; Schell et al. 2017; Segaert et al. 2018; Zhang and Pylkkänen 2018). Furthermore, we followed the approach of the MEG binding literature of not analysing the first word when it is not possible to match it between conditions (e.g., Bemis and Pylkkänen 2011; Segaert et al. 2018).

Behavioural evidence from previous use of this paradigm has shown that participants judge pseudo-verb sentences with the correct morphological inflection to be valid sentences (e.g., “*she grushes*”), but judge sentences with the incorrect inflection (e.g., “*she grush*”, “*I grushes*”) and pseudo-verb wordlists (e.g., “*cugged grushes*”) to be invalid sentences (Poulisse et al. 2019; Poulisse et al. 2020). This is evidence that listeners do not attempt to bind the two pseudo-verbs together when in a wordlist as, if they did, they would judge it to be a valid sentence (e.g., if they had interpreted “*cugged grushes*” as a deverbal adjective and noun pairing). We therefore consider it highly likely that participants in our study were engaging in syntactic binding when a minimal sentence was presented, but not when a wordlist was presented (in line with the assumptions of other studies of minimal binding; Bemis and Pylkkänen 2011, 2013; Segaert et al. 2018; Blanco-Elorrieta et al. 2018; Poulisse et al. 2020).^1^

To construct the experimental items, we used a set of 20 pseudo-verbs created by Ullman et al. (1997).^2^ All pseudo-verbs were monosyllabic and could be inflected according to the grammatical rules of regular English verbs. We combined each pseudo-verb with three different morphological affixes (no affix; +s; +ed) to create 60 possible pseudo-verb-affix combinations. In English, only certain pronouns may be combined with certain affixes (e.g., “*she grushes*” is acceptable, but “*I grushes*” is not). Using a list of six pronouns (I, you, he, she, they, we), we created 120 sentence items by pairing each pseudo-verb-affix with two different pronouns that were syntactically appropriate for the corresponding affix, such that syntactic binding may plausibly occur (e.g., “*I dotch*”, “*she grushes*”, “*they cugged*”). To create the wordlist items, we paired together two different pseudo-verb-affix stimuli for which no syntactic binding occurred (e.g., “*cugged grushes*”, “*dotch traffed*”). Each pseudo-verb-affix stimulus occurred twice as the first word in a pair and twice as the second word in a pair, creating a total of 120 wordlist items. We ensured that the two words within each wordlist pair always consisted of a different pseudo-verb and a different affix.

We also created 120 filler items. Sixty of the fillers consisted of reversed speech and were included as a detection task for the participants. We reversed the speech of each of the 60 pseudo-verb-affix combinations and then paired each with either a non-reversed pseudo-verb-affix or pronoun (half as the first word of the pair and half as the second word). A further 60 filler items were used to increase the variability of stimuli presented to participants (Roland et al. 2007). Thirty such items consisted of two pronouns (e.g., “*she I*”); this contrasted the experimental sentence items in which a pronoun was followed by a pseudo-verb (e.g., “*she grushes*”). The other 30 items consisted of a pseudo-verb-affix stimulus followed by one of five possible adverbs (early, promptly, quickly, rarely, safely; e.g., “*cugged quickly*”); this contrasted the experimental wordlist items in which a pseudo-verb was followed by another pseudo-verb (e.g., “*cugged grushes*”). Overall, this set of fillers created a task and variability for the participants – the filler items did not act as a control to the experimental items and were not analysed.

All auditory stimuli were spoken by a native English male speaker and normalized to 1db volume. Each experimental item consisted of two separate audio files for word 1 and word 2. Within each audio file, the onset of the word began at exactly 0s – we achieved this using the audio-editing software *Praat* (Boersma 2001).

### Experimental Procedure

The participants’ task was to detect the reversed speech (which only occurred on filler trials). On each trial, participants were auditorily presented with a two-word phrase (**Figure 1**). At the beginning of each trial, a fixation cross was presented on screen for 1.2s with no audio – this represented our baseline period. At 1.2s, the onset of the first word auditory presentation started. The length of word 1 audio varied between 0.3s and 0.6s. There was then a brief pause before the onset of word 2 audio presentation began, exactly 1.2s after the word 1 onset (i.e., at 2.4s in the overall trial timings). This time interval means that the lexical processing of word 1 (which occurs within the first 0.6 seconds of auditory onset; MacGregor et al. 2012) will have been completed before the onset of word 2. Again, the length of the word 2 audio varied between 0.3s and 0.6s. The fixation crossed remained on screen throughout the word 1 and word 2 presentation to limit participants’ eye movements.

**Figure 1.**
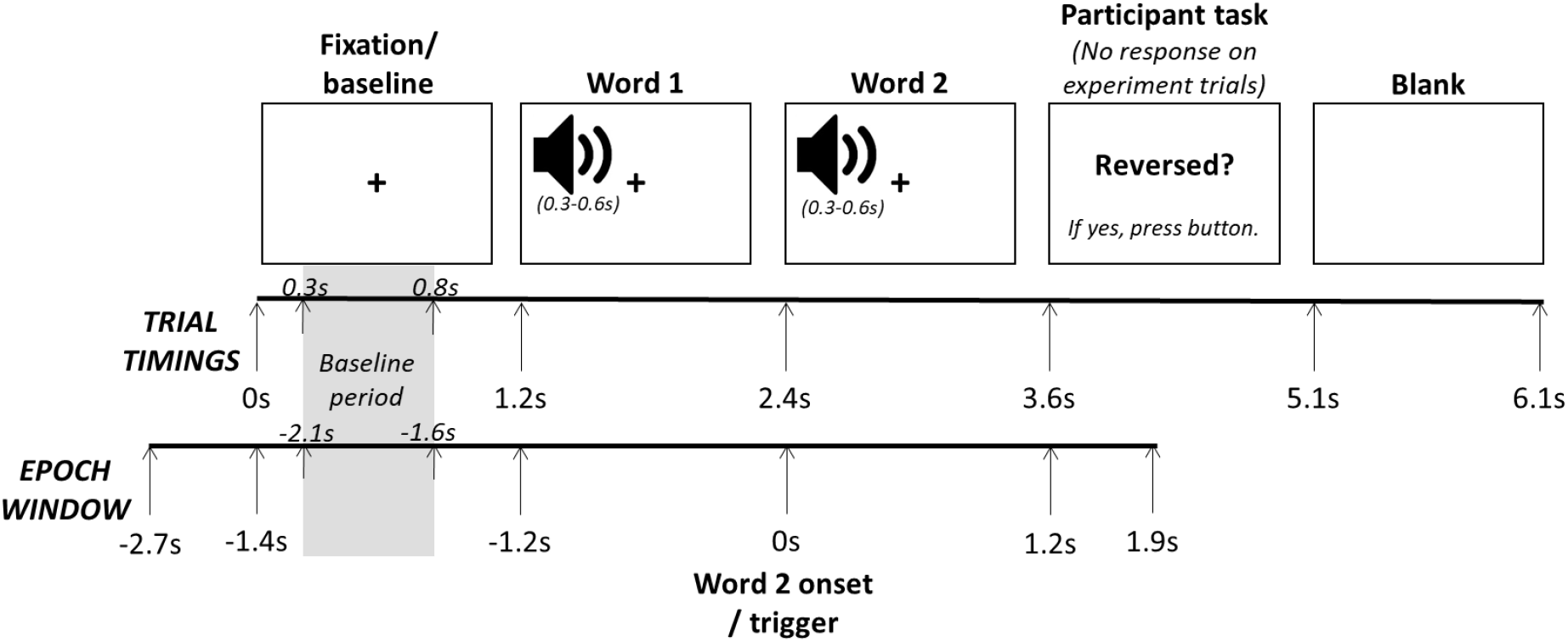
Stimuli presentation timings per trial and the related epoch window. The trial timing shown in the figure relates to the onset of the stimuli. The length of the audio files of word 1 and word 2 varied between 0.3s and 0.6s. Stimuli presentation and trigger signals were controlled using E-Prime (Schneider et al. 2002). Visual stimuli were presented using a PROPixx projector, and auditory stimuli were presented using the *Elekta* audio system and MEG-compatible ear phones. Participants’ motor responses were recorded using a NAtA button pad.

Following the word 2 audio presentation, there was a brief pause until at exactly 1.2s after the word 2 onset (i.e., at 3.6s in the overall trial timings), the phrase “Reversed?” appeared on the screen – this was the participant’s decision task. Participants were instructed to press a button if part of the speech was reversed (half of the participants used their left index finger, and half used their right index finger), but to do nothing if the speech was not reversed. There was no difference in response decision processes (i.e., no button press) between the critical experimental conditions of interest (sentence vs. wordlist). The “Reversed?” question remained on screen for 1.5s (in which time the participant pressed a button or did nothing). Following this, the screen went blank for 1s before the next trial began with the presentation of the fixation cross. In total, each trial lasted 6.1s. Each participant completed 360 trials (consisting of 240 experimental trials and 120 filler trials) in a unique randomized order, divided into six blocks of 60 trials each. Before beginning the task, participants completed 23 practice trials that were similar to the experimental and filler items used in the main task.

As expected, participants were highly accurate at detecting the reverse speech on the filler trials (*M* = 94.6%, *SD* = 2.3%, *Range* = 82-98%), indicating that they were closely listening to the filler and experimental stimuli throughout (since they did not know when the reversed speech would be presented).

### Data Acquisition

During the task, ongoing MEG data were recorded using the TRIUX™ system from *Elekta* (Elekta AB, Stockholm, Sweden). This system has 102 magnetometers and 204 planar gradiometers. These are placed at 306 locations, each having one magnetometer and a set of two orthogonal gradiometers. The data were collected using a sampling rate of 1000 Hz and was stored for offline analyses. Prior to sampling, a lowpass filter of ~250 Hz was applied. Four head position indicator coils (HPIs) were placed behind the left and right ear, as well as on the left and right forehead just below the hairline. The positions of the HPIs, the nasion, the left and right preauricular points, as well as the surface points of the scalp, were digitized using a *Polhemus*™ 3D device to facilitate later co-registration with anatomical brain scans. Additional electrooculography (EOG) and electrocardiogram (ECG) data were collected using methods compatible with the TRIUX™ system.

Anatomical high-resolution T1 brain images were acquired for participants at a later session using a MAGNETOM Prisma 3T MRI system from *Siemens* (Siemens Healthcare, Erlangen, Germany). These images were used for the reconstruction of individual head shapes to create forward models for the source localization analyses.

#### MEG Pre-processing

The offline processing and analyses of the data were performed using functions from the Fieldtrip software package (Oostenveld et al. 2011) and custom scripts in the MATLAB environment. First, we applied a 0.1 Hz high-pass filter to the MEG data to remove slow frequency drift in the data. The data were segmented into epochs aligned to the onset of the auditory presentation of the second word from −2.7s to 1.9s (see **Figure 1**) and demeaned. We applied a baseline-subtraction in the time domain to the data (i.e., each trial was subtracted by the mean activity −0.1 to 0s prior to the onset of the fixation). We ran automatic artefact rejection routine implemented in Fieldtrip to remove superconducting quantum interference device (SQUID) jumps.

We first removed all filler trials as we were specifically interested in the difference in neural responses between the experimental sentence and wordlist conditions (i.e., our contrast of interest). We further removed experimental trials for which the participant incorrectly responded with a button press (i.e., indicated that the speech was reversed when it was not) and experimental trials during which the participant made an accidental button press before the response screen (i.e., during the fixation cross and/or auditory presentation). We then visually inspected the waveforms of each experimental trial and removed trials that contained excessive signal artefacts (e.g., large sensor jumps or gross motor movement by the participant); on average, we removed 16% of experimental trials (38.3/240) per participant (*Range* = 4-47%). Following all this, there was an average of 101 sentence trials (*SD* = 12.2) and 100 wordlist trials (*SD* = 11.6) per participant that were usable for the analyses (out of a maximal 120 trials per experimental condition).

We further removed any persistently poor channels that contained excessive noise or flatlined (*Mean* channels removed per participant = 2.46; *SD* = 2.86; *Range* = 0-9). We then used a spline interpolation weighted neighbourhood estimate to interpolate across the removed channels per participant. Ocular and cardiac artefacts were removed from the data using an independent component analysis (ICA) (*Mean* artefacts removed per participant = 1.75; *SD* = 0.68; *Range* = 1-3). We identified these components from their stereotypical topography and time course, as well as by comparisons with the recorded ECG and EOG time courses.

### Statistical Analyses

#### Time-frequency

For the frequency range 1-30 Hz (1 Hz steps), we obtained time-frequency representations (TFRs) of power for each trial using sliding Hanning tapers with an adaptive time window of three cycles for each frequency (ΔT = 3/f). This approach has also been used in a number of previous studies (e.g., Whitmarsh et al. 2011; Van Diepen and Mazaheri 2017; Segaert et al. 2018). For each participant, the data for the planar gradiometer pairs was added to create a 102-channel combined planar map in sensor space, and we baseline-corrected the data using the oscillatory activity during the fixation cross presented at the beginning of the trial. Specifically, using the ‘absolute’ baseline parameter in Fieldtrip, we subtracted the mean of the power of each frequency in a 0.5s period of the fixation cross presentation (that being −2.1s to −1.6s in the epoch window) from all other power values. We calculated the TFRs separately per experimental condition for each participant and then averaged across all participants.

We assessed the statistical differences in time-frequency power between the sentence and wordlist conditions across participants using a cluster-level randomization test (incorporated in the Fieldtrip software), which circumvents the type-1 error rate in a situation involving multiple comparisons (i.e., multiple channels and time-frequency points; Maris and Oostenveld 2007). This approach first clusters the data in sensor space depending on whether the contrast between the two conditions exceeds a dependent samples *t*-test threshold of *p* < .05 (two-tailed). In line with Segaert et al. (2018), we used the following pre-defined frequency bands: theta (4-7 Hz), alpha (8-12 Hz), low beta (15-20 Hz) and high beta (25-30 Hz). We considered a cluster to consist of at least two significant adjacent combined planar gradiometers. A Monte Carlo *p*-value of a cluster was then obtained by calculating the number of times the *t*-statistics in the shuffled distribution is higher than the original *t*-statistic obtained when contrasting conditions across 1000 random permutations. We first performed the analyses within the time window of interest, centred around the presentation of the second word (−0.5s to 1s of the epoch), as this is where we expect to observe time-frequency effects of syntax composition. We then performed the analyses across the complete timeframe of the auditory presentation (−1.3s to 1.2s of the epoch).

To ensure that the observed alpha power changes were not just the spectral representation of the event-related fields (ERFs), the ERF components were subtracted from the TFR (Mazaheri and Picton 2005). The subtraction was achieved by first generating the time frequency decomposition of the ERF data for each condition and participant separately. Next, the time frequency power spectra of the ERF were subtracted from the time frequency power spectra of the MEG signal (not baseline corrected) for each condition. The subsequent power changes in the time-frequency domain were used to generate time frequency power spectra differences between experimental conditions (sentence vs. wordlist). We then reanalysed the ERF-adjusted TRFs using the same statistical methods outlined above.

#### Source Localization

A realistically shaped description of each participant’s brain was constructed using individual head models obtained using the Polhemus 3D digitizer and the acquired MRI anatomical brain scan (where available). Specifically, using the Iterative Closest Point (ICP) algorithm (Besl and McKay 1992) implemented in Fieldtrip, we manually aligned the MRI images to the digitized scalp surface of each individual. The alignment according to the MEG sensor array was done relative to four digitized head position indicator coils. The MRI images were then segmented in Fieldtrip and a realistically shaped single-shell description of the brain-skull interface was constructed (Nolte 2003).

Source estimation of the time-locked MEG data was performed using a frequency-domain beam-forming approach (dynamic imaging of coherent sources [DICS]; Gross et al. 2001), which uses adaptive spatial filters to localize power in the entire brain. Specifically, we obtained the cross-spectra density (CSD) matrices in each condition; the CSDs were calculated using Fourier transform in combination with multi-tapers (+/- 3 Hz smoothing). Based on our sensor results, we focused our analysis on 10 Hz (7-13 Hz) activity centred around −0.15 to 0.15s surrounding the onset of the second word.^3^ Our regularization parameter was set to 5%.

The brain volume of each individual participant was discretized to a grid with a 0.8cm resolution and the lead field was calculated for each grid point. A common filter was calculated for the sentence and wordlist condition and then applied for the data separately for the individual conditions (Whitmarsh et al. 2011; Mazaheri et al. 2014). The source estimates of the individual participants’ functional data along with the individual anatomical MRI images were warped into a Montreal Neurological Institute (MNI) standard brain (Quebec, Canada; http://www.bic.mni.mcgill.ca/brainweb) before averaging and statistics.

We performed cluster-based randomization tests to identify the grid-points in which there was a significant difference in power between the two experimental conditions, guided by the significant findings of the time-frequency analyses (i.e., alpha activity surrounding the onset of the second word) as is a common approach within the field (Wang et al. 2012, 2017; Magazzini et al. 2016). We followed the approach taken in previous MEG literature by directly comparing the two conditions without a baseline correction (Mazaheri et al. 2009; Magazzini et al. 2016; Wang et al. 2017). We used these clusters to identify specific regions of interest (ROIs) of the condition difference. We derived anatomical labels of these regions from Brodmann’s map and from the Automated Anatomical Labelling Atlas (Tzourio-Mazoyer et al. 2002).

#### Inter-regional Connectivity

We performed inter-regional connectivity analyses on four distinct ROIs identified as showing a significant condition difference in the source localization analyses. TFRs of the complex Fourier spectra for each trial at each ROI was obtained by using a sliding Hanning tapers with an adaptive time window of three cycles for each frequency (ΔT = 3/f). We calculated the inter-regional phase-locked indices of the oscillatory activity (2-30 Hz) between the four ROIs (creating six inter-regional connections) for the time period of interest (−0.5s to 1s of the epoch) separately for the sentence condition and wordlist condition (following Lachaux et al. 1999; Bastos and Schoffelen 2016). The phase locking index (PLI) is calculated across all trials for each condition, frequency and time point. We used the following formula to calculate the PLI:

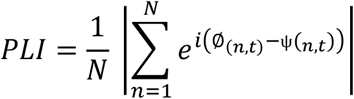

Applied to our study, the PLI reflects the consistency of the phase difference of oscillatory activity across trials between two ROIs. Here: *N* is the number of trials; Ø_(n,t)_ is the phase (obtained from the complex spectra) at time *t* in trial *n* in one ROI; and ψ_(n,t)_ is the phase at time *t* in trial *n* in the other ROI. A PLI of 0 indicates no phase locking between the activities of the two ROIs, whereas a PLI of 1 is indicative of perfect phase locking.

We then performed cluster-level randomization tests, involving 1000 permutations, (Maris and Oostenveld 2007) on the PLIs in order to identify the inter-regional connections in which there was a significant difference in phase-locking between the two experimental conditions (for a similar statistical approach see, Schmidt et al. 2014). The use of a cluster-level randomization test, in which a Monte Carlo *p*-value is obtained, enables the control of multiple comparisons of different ROIs and time-frequency points. To correct for the multiple analyses performed across the six different inter-regional connections, we applied a Bonferroni correction (α / *n* comparisons) to our critical *p* value of interest (.05/6 = .0083). We used the same pre-defined frequency bands as for the oscillatory time-frequency analyses: theta (4-7 Hz); alpha (8-12 Hz); low beta (15-20 Hz) and high beta (25-30 Hz).

## Results

We compared participants’ MEG activity during the comprehension of minimal sentences that were highly likely to require binding (a pronoun combined with a pseudo-verb with the corresponding morphological inflection; “*she grushes*”) to wordlists that did not require binding (two pseudo-verbs; “*cugged grushes*”). Our findings reveal two key mechanisms of syntactic binding.

### Less alpha power in the left-lateralized brain network when syntactic binding occurs

The grand-average of the time-frequency representations (TFR) of power averaged across all sensors aligned to the onset of the second word are summarized in **Figure 2A** for the sentence condition in which syntactic binding was highly likely to occur, and the wordlist condition in which binding was highly unlikely.^4^ In both conditions, there are power increases in alpha and low beta surrounding the presentation of the second word (at 0s; “*grushes*”). Approximately 0.5s after the second word presentation, the strength of the alpha and low beta power signal becomes less pronounced in both conditions (although it is still positive compared to baseline).

The statistical results of the cluster-based permutation tests (which controlled for multiple comparisons of different channels and time-frequency points) of the time window of interest revealed that there was a significant condition difference in the alpha frequency range (8-12 Hz) surrounding the presentation of the second word where alpha power was lower in the sentence condition, compared to the wordlist condition (*p* = .021, see **Figure 2B-C**). This difference was maximal around the onset of the second word (−0.05s to 0.1) over a cluster of sensors predominantly in the left-frontal region (**Figure 2B**). Importantly, our observed oscillatory effect of alpha was distinct from the evoked fields as, when we analysed the ERF-adjusted TFRs, we found the same significant alpha power cluster at −0.05s to 0.1 (*p* = .022); see **Supplementary Materials 2** for additional figures. We found no significant difference in oscillatory activity within the other analysed frequency bands: theta (4-7 Hz), low beta (15-20 Hz) and high beta (25-30 Hz). When we analysed the complete timeframe of the auditory stimuli presentation, the significant condition difference in alpha surrounding the second word remained significant (−0.05s to 0.1s, *p* = .024), while we observed no other significant effects at any other time points or frequencies.

**Figure 2.**
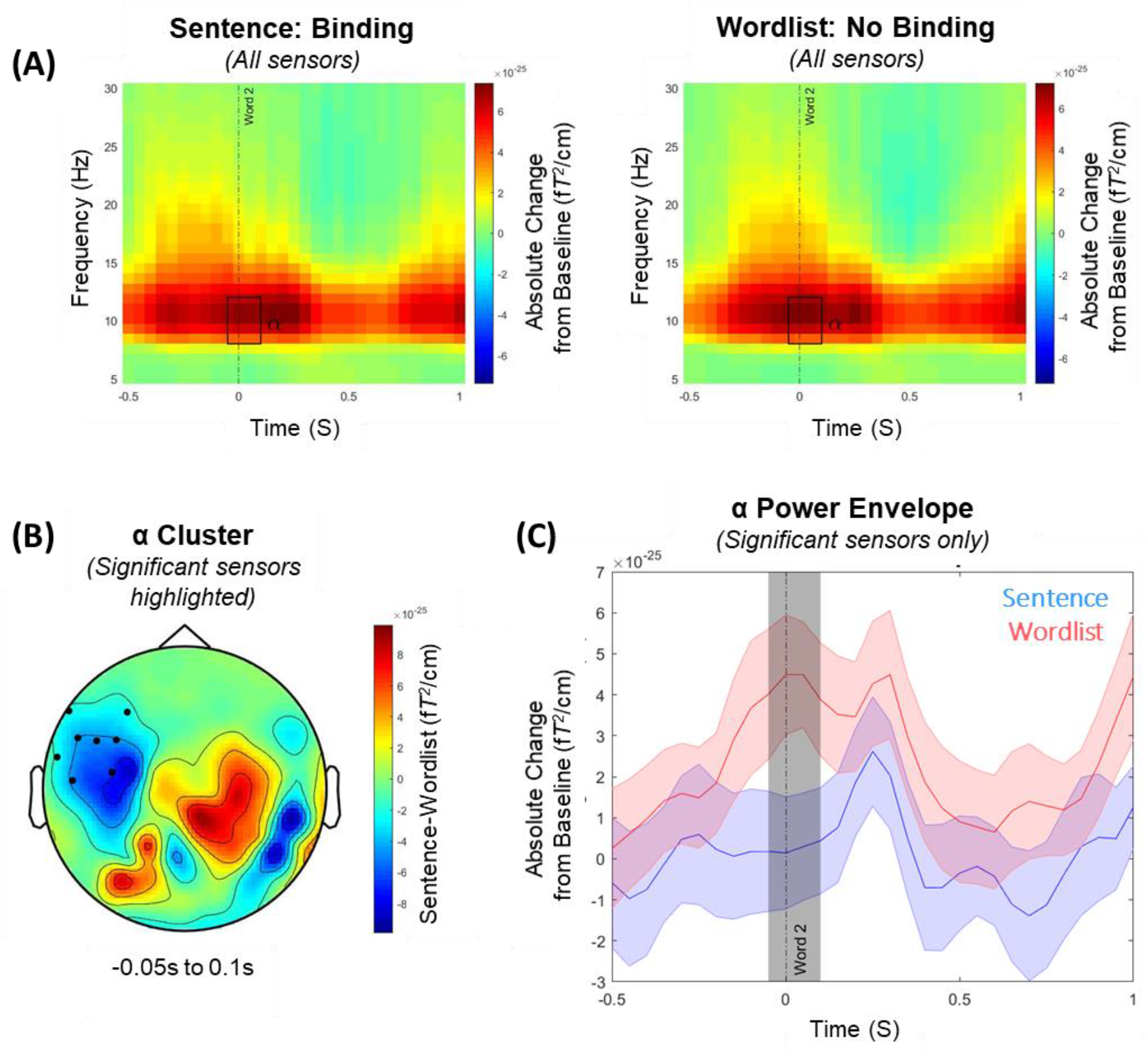
**(A)** Time-frequency representations of power averaged across all sensors, expressed as an absolute change from the baseline period (i.e., −2.1s to −1.6s before the onset of the second word) for the sentence condition [left panel] in which syntactic binding was highly likely to occur (e.g., “*she grushes*”), and the wordlist condition [right panel] in which binding was highly unlikely (e.g., “*cugged grushes*”). Time relates to the main time period of interest aligned to the onset of the second word (at 0s). The rectangle highlights the time period where we observed the significant difference in alpha power (8-12 Hz) between the two conditions (−0.05s to 0.1s; *p* = .021). **(B)** The scalp topography of the condition contrast (sentence minus wordlist) of the averaged alpha power activity in the time window (−0.05s to 0.1s) where we observed the significant difference in alpha power between the two conditions. The black dots illustrate where this effect was largest at the scalp level. **(C)** The time course of the alpha power envelope for the sensors showing a significant difference in power between the Sentence (blue) and Wordlist (red) conditions. The shaded coloured areas represent the standard error of the mean. The shaded grey area indicates the time window in which the difference between conditions is significant, centred around the presentation of the second word (−0.05s to 0.1s).

Source analyses co-registered on the participants’ anatomical MRI brain scans indicated that the significant condition difference in alpha power (8-12 Hz, −0.05s to 0.1s) was localized to a network of left-lateralized brain regions which are typically associated with language function (**Figure 3A**). Within this network, we identified four brain areas (see **Figure 3B**) in which significant differences were found between the sentence condition and wordlist condition: the left inferior frontal gyrus (BA44; peak coordinates [-42 7 17]); the left angular gyrus (BA39; peak coordinates [−58 −48 29]); the left middle/inferior temporal cortex (BA21; peak coordinates [−60 −30 −18]); and the left anterior frontal gyrus (BA46; peak coordinates [-40 43 7]).

**Figure 3.**
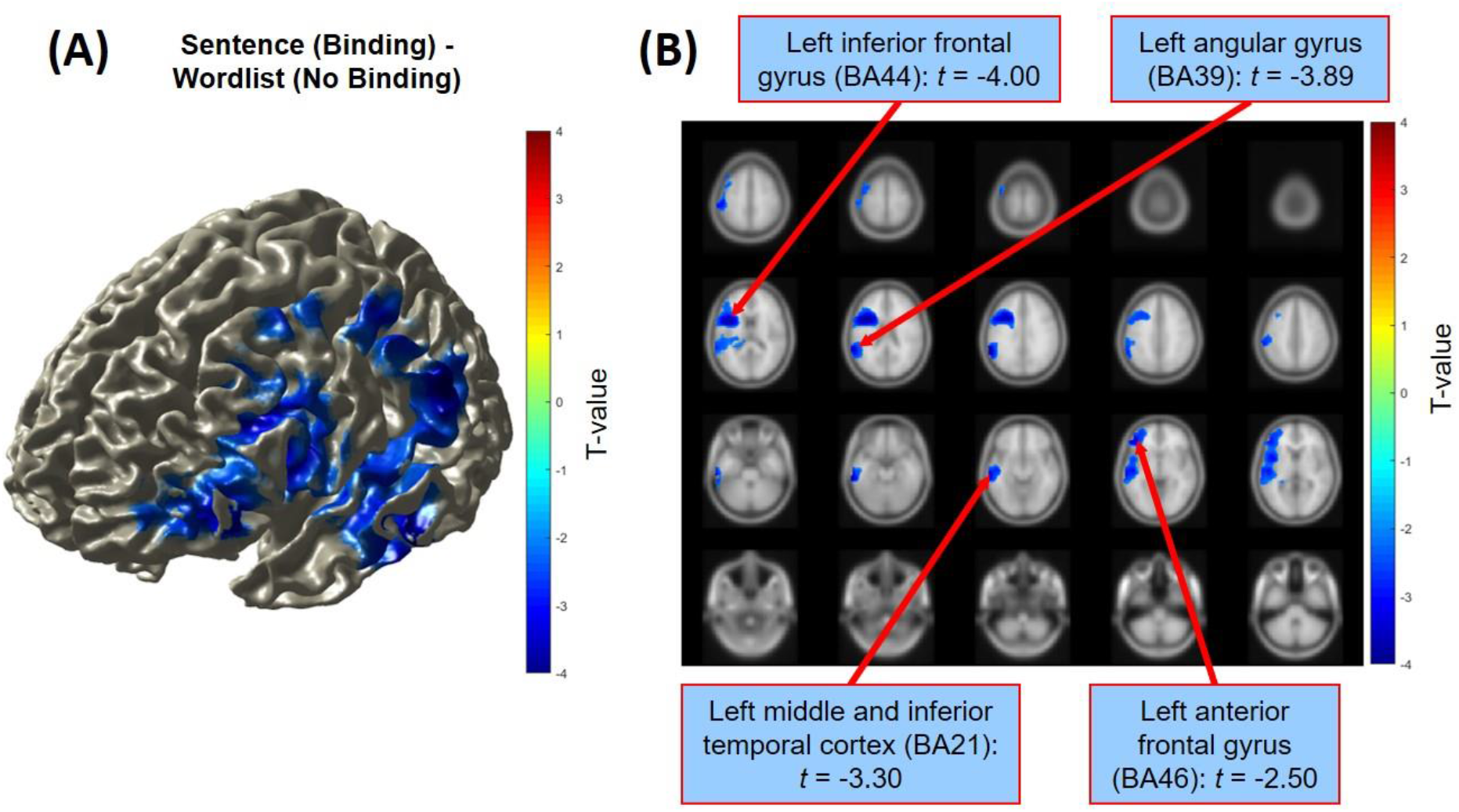
Source localization estimates of the condition difference in alpha power (8-12 Hz) of the sentence condition (e.g., “*she grushes*”) minus the wordlist condition (e.g., “*cugged grushes*”) surrounding the presentation of the target word (−0.15s to 0.15s) as shown for a surface **(A)** and sliced **(B)** view of the brain. The displays are masked for significant clusters only (*p* < .05). The condition difference was maximal over the left-frontal areas of the brain, with significant differences observed in the left inferior frontal gyrus, the left angular gyrus, the left middle/inferior temporal cortex, and the left anterior frontal gyrus.

### During syntactic binding, there is decreased alpha phase-locking between the left inferior frontal gyrus and the left middle/inferior temporal cortex

We calculated the inter-regional phase-locked indexes between the peak coordinates of the four brain areas identified as displaying significant condition differences in the source localization analyses (**Figure 4A**). The statistical results of the cluster-based permutation tests (which controlled for multiple comparisons of different ROIs and time-frequency points) revealed a significant condition difference (*p* = .005) in inter-regional phase-locking in alpha activity (8-12 Hz) between the left inferior frontal gyrus (BA44) and the left middle/inferior temporal cortex (BA21) following the presentation of the second word that required binding (around 0.15s to 0.25s). During this time period, there was significantly less alpha phase-locking between these two brain regions in the sentence condition, compared to the wordlist condition (**Figures 4B-D**). While we are aware that amplitude may correlate with PLI (Van Diepen and Mazaheri 2018), we consider that it is unlikely that our observed condition difference in the PLIs is driven by amplitude differences alone because it occurred at a time interval when there was *not* the greatest amplitude difference between the two conditions, and because we only observed the effect between two specific ROIs, not the whole analysed left-lateralized brain network. We did not find any other significant condition differences in inter-regional phase-locking in the other connections between our ROIs (**Figure 4E**).

**Figure 4.**
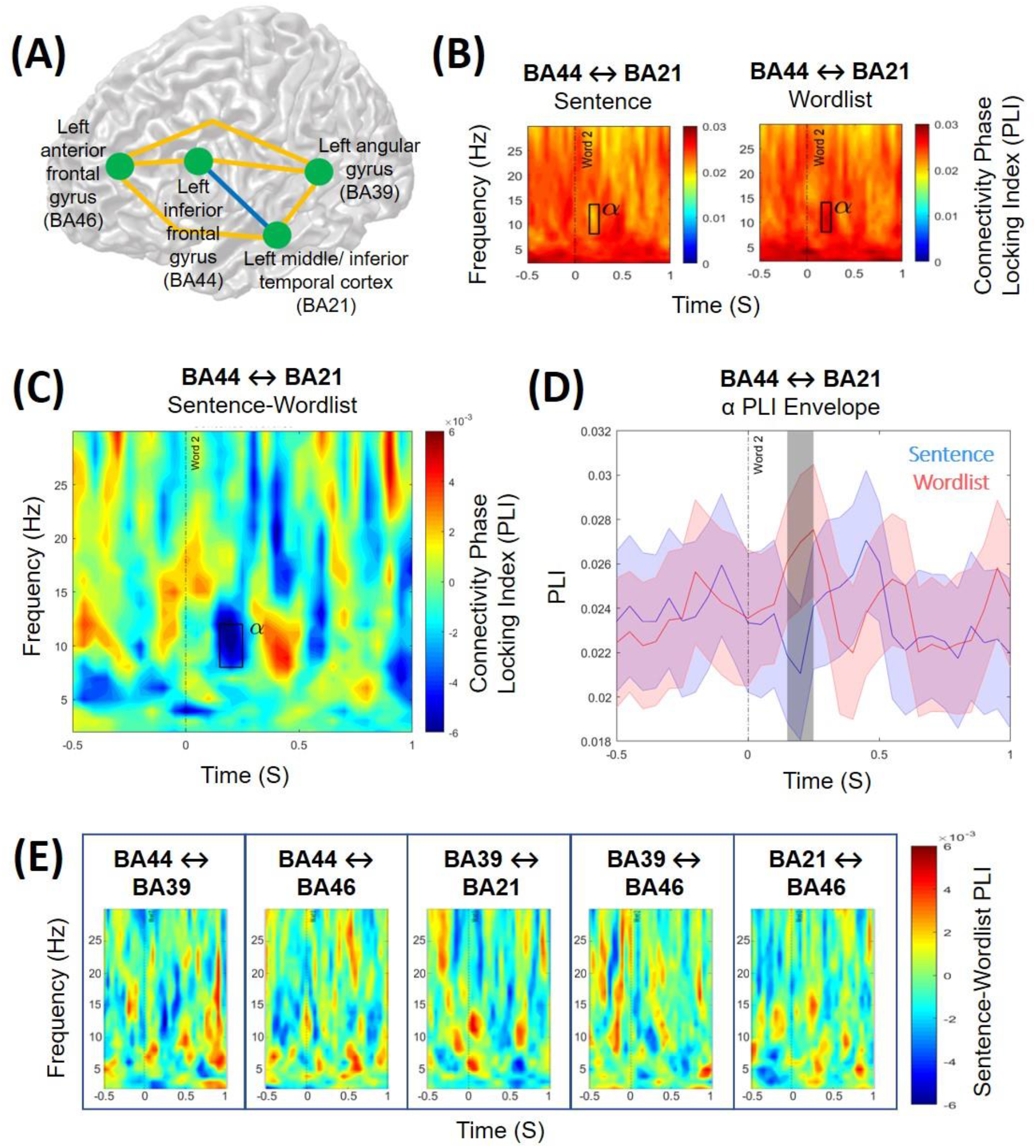
**(A)** Inter-regional phase-locking differences of oscillatory activity (2-30 Hz) were calculated between four distinct brain regions of interest in the left hemisphere for the time period of interest centred around the presentation of the target word (−0.5s to 1s). **(B)** A significant condition difference in inter-regional phase-locking was observed between the left interior frontal gyrus (BA44) and the left middle and inferior temporal cortex (BA21) in the alpha frequency range (8-12 Hz). Time-frequency estimates of the phase-locking index are shown for the Sentence condition (“*she grushes*”) and the Wordlist condition (“*cugged grushes*”). There was significantly greater phase-locking of alpha activity between BA44 and BA21 in the Wordlist condition, compared to the Sentence condition, 0.15s to 0.25s following the presentation of the target word (*p* = .005), as highlighted by the rectangle. **(C)** The condition contrast (Sentence minus Wordlist) of the phase-locking interest between BA44 and BA21. The rectangle highlights the time window and frequency range in which significantly different phase-locking was observed between the conditions. **(D)** The time course of the phase-locking index for the alpha frequency range (8-12 Hz) between BA44 and BA21 for the Sentence condition (red) and Wordlist condition (blue). The shaded coloured areas represent the standard error of the mean. The shaded grey area indicates the time window in which the difference between conditions was significant (0.15s to 0.25s). **(E)** The inter-regional phase-locking index of the condition contrast (Sentence minus Wordlist) between the other brain regions of interest; no significant differences were found between the Sentence and Wordlist conditions.

## Discussion

The findings of our MEG study suggest that minimal syntax composition is associated with distinct oscillatory changes in alpha band activity and the engagement of the left-lateralized language network of the brain. In the reported experiment we compared minimal pseudo-verb sentences for which it was highly likely that syntactic binding may plausibly occur (e.g., “*she grushes*”) to wordlists for which binding was highly unlikely to occur (e.g., “*cugged grushes*”). We found that surrounding (−0.05s to 0.1s) the presentation of the target word (“*grushes*”), alpha power was significantly less in the sentence, compared to the wordlist, condition. The sources of this condition difference were localized to left-lateralized brain regions, including the inferior frontal gyrus (IFG), angular gyrus, middle/inferior temporal cortex, and anterior frontal gyrus. Moreover, following the presentation of the target word (0.15s to 0.25s), we observed decreased alpha phase-locking between the LIFG and the left middle/inferior temporal cortex in the sentence, compared to the wordlist, condition, indicating that syntactic binding is associated with less coupling of alpha activity between these regions.

### Modulation in alpha power in a network of left-lateralized language regions reflects an expectation of binding to occur

Compared to baseline, we observed an increase in alpha power in both conditions during the comprehension of the second word, but critically this power increase was *less* when syntactic binding was required, compared to when no binding was required. This modulation in alpha power is consistent with existing evidence of alpha oscillatory activity in compositional processing (Lam et al. 2016; Segaert et al. 2018; Gastaldon et al. 2020) and the view that neural oscillations subserve the processing of syntactic information within language-relevant cortices (Meyer 2018; Prystauka and Lewis 2019). Our findings further demonstrate that alpha operates in auditory language processing in addition to its role in visual processing in occipital areas (Mazaheri et al. 2014; Zumer et al. 2014), supporting a varied functionality of alpha oscillations in multiple sensory systems in different cortical regions (Foxe and Snyder 2011).

We suggest that, in our experiment, the observed lesser alpha power in the binding context around the presentation of the target word reflects an expectation of binding to occur. When comprehending linguistic input, we build expectations in order to predict upcoming words (Chang et al. 2006; Kuperberg and Jaeger 2016); indeed, probability estimates are considered to be inherent within the neural linguistic system (Kutas et al. 2011). Thus, if the first word was a pronoun, as opposed to a pseudo-verb, the participant may reasonably expect that binding was likely to be required given their existing knowledge (based on language use in everyday life) about the properties and syntactic function of pronouns. This interpretation is consistent with studies that have found greater alpha suppression (i.e., decreased alpha power compared to baseline) when participants comprehend a highly predictive, compared to a less predictive, sentence (Piai et al. 2014; Rommers et al. 2017; Wang et al. 2017) and the proposed role of alpha power decreases in controlling the allocation of the brain’s resources (Jensen and Mazaheri 2010; Klimesch 2012). In particular, decreased alpha power limits cortical excitability, thereby allowing for continual processing regardless of alpha phase (medium-to-high attentional state); whereas, when alpha power is higher, cortical processing is more discontinuous as it depends on the phase of the alpha rhythm (rhythmic attentional state) (Jensen et al. 2014; Van Diepen et al. 2019; Alavash et al. 2021). Our finding of less alpha power around the presentation of the target word that required binding (compared to a no binding context) may therefore reflect the initiation of anticipatory binding processes (for which a medium-to-high attentional state is required), along with the increased engagement of brain regions involved in syntactic binding.

The observed condition difference in alpha power was localized to a left-lateralized network of brain regions, consistent with established neurobiological models of linguistic processing (Hagoort 2003; Tyler and Marslen-Wilson 2008; Friederici 2011). The implication of the LIFG (BA44) is expected given the region’s proposed role in managing the combination of words into a coherent syntactic structure (Hagoort 2005; Snijders et al. 2008; Segaert et al. 2012; Uddén et al. 2019) and in computing dependency structures (Lopopolo et al. 2021). Indeed, the LIFG has been identified in previous studies of minimal syntactic binding (Zaccarella and Friederici 2015; Zaccarella et al. 2017) and top-down predictive processing (Matchin et al. 2017; Strijkers et al. 2019). The other three left-lateralized regions we identified - angular gyrus (BA39), anterior frontal gyrus (BA46), and middle/inferior temporal cortex (BA21) - have also been found to contribute towards syntactic processing (Giraud 2004; Humphries et al. 2006; Menenti et al. 2011; Segaert et al. 2012; Matchin et al. 2017). Our findings therefore suggest that successful composition of the syntactic properties between words in a sentence is driven by the engagement of a distributed network of left-lateralized regions (including the frontal gyri, temporal cortex and angular gyrus), and critically their coordination (as we discuss in more detail in the next section). Moreover, given that we did not observe any effect of compositionality in the anterior temporal lobe (ATL) in our task (in which contributions from semantics were minimized), our findings further add to the current evidence that the functional role of the ATL relates primarily to semantic, not syntactic, composition (Del Prato and Pylkkänen 2014; Wilson et al. 2014; Kim and Pylkkänen 2019; Pylkkänen 2019).

However, our findings of alpha power decrease is somewhat at odds with Segaert et al. (2018) who, using a similar paradigm but with EEG, found syntactic binding to be associated with alpha power increases. One explanation for this difference may reflect MEG vs. EEG differences in spatial coherence and sensitivity to deeper brain tissues that can lead to differences in detectable power (Lopes da Silva 2013; Bénar et al. 2019). In particular, MEG and EEG differ in their sensitivity to the radical and tangential components of the dipolar sources in the brain, which can lead to the two producing diverging estimates for the same cognitive process (Edgar et al. 2003; Dehghani et al. 2010; Fonteneau et al. 2015; Ross et al. 2020). An alternative explanation for the study differences may relate to morpho-syntactic differences between English (used in this study) and Dutch (used in Segaert et al. 2018). Compared to English, Dutch is more morphologically complex in that there are a greater number of possible verb inflections (to signal its binding to a preceding pronoun) and there are more rules on inflection usage (Booij 2019). This could mean that Dutch listeners focus more on morphological congruency, whereas English speakers focus more on overall sentential composition, leading to group differences in detectable alpha power. Further work is therefore needed to uncover how differences in the complexity of the inflection system of a language may drive differences in the oscillatory signature of syntax composition even at its most basic two-word level. Nevertheless, the findings of our study and Segaert et al. (2018) should not be considered as directly opposed, but instead reflective of the different components of the wider neural processes involved in syntax composition (the intricacies of which may differ between languages or which may be differently detected by MEG vs. EEG), and the more general functionality of alpha oscillations in syntax processing (Meyer 2018; Prystauka and Lewis 2019).

### Decreased alpha band phase-locking in the language network reflects strengthened network communication required for successful syntactic binding

Inter-regional connectivity analyses revealed that syntactic binding was associated with less coupling of alpha between the LIFG and the left middle/inferior temporal cortex: we found *less* alpha phase-locking in the sentence, compared to the wordlist, condition following the presentation of the target word (0.15s to 0.25s). The time window of the effect, after the auditory processing of the target word (which typically occurs within the first 0.1s; Zouridakis et al. 1998), suggests that it reflects the underlying syntax composition mechanisms taking place as opposed to an expectation of binding to occur. Our finding is consistent with evidence that less alpha band coupling (also referred to as alpha desynchronization) between relevant brain regions is beneficial for language comprehension and can predict successful sentence encoding (Becker et al. 2013; Magazzini et al. 2016; Vassileiou et al. 2018; Alavash et al. 2021). This functional connectivity reflects the dynamic interaction among distributed brain regions that are subserved by deep white matter pathways and which communicate through frequency-specific networks (Schoffelen et al. 2017; White et al. 2018; Sarubbo et al. 2020). This communication may operate through top-down mechanisms of cortical disinhibition, such that alpha band desynchronization serves to functionally disinhibit the cortex to enable information to be transferred from and to specific areas for processing (Jensen and Mazaheri 2010; Klimesch 2012; Sadaghiani and Kleinschmidt 2016).

We suggest that similar mechanisms of frequency-specific communication and alpha-driven disinhibition were operating in our task when participants comprehended sentences that needed binding (“*she grushes*”). In order to bind the two words together into a minimal syntactic structure, increased information, such as the labels of syntactic components (Murphy 2015), needed to be transferred between the LIFG and the middle/inferior temporal cortex. Thus, the less alpha band synchronization between these two regions, the greater the cortical disinhibition; this, in turn, strengthens the communication within the cortical and oscillatory network involved in syntax composition, thereby enhancing participants’ processing.

Literature suggests that an increase in alpha activity in a brain area reflects gating of information in that area (Klimesch et al. 2007; Jensen and Mazaheri 2010; Scheeringa et al. 2011; Van Diepen et al. 2019). The precise mechanism underlying this inhibition has not yet been fully elucidated, though one proposition is that an alpha cycle may reflect pulses that could be inhibiting the firing rate of neurons (Haegens et al. 2011). We ourselves have previously speculated that alpha activity can be viewed as rhythmic pulses of inhibition and excitability that cycle on and off approximately every 100ms (Mazaheri and Jensen 2010). Within a region, alpha activity can be seen as serving as ‘gain-control’ by limiting the duty-cycle (i.e., the ratio between on/off) of information processing in a cortical region (Spaak et al. 2012). Based on Van Diepen et al.’s (2019) recent model of alpha synchronization, we speculate that in a situation where alpha activity is synchronous between two brain areas, information flow between them is gated because of the restriction posed by the duty cycle of processing; however, a suppression of alpha activity (i.e., decrease in synchrony between brain areas) would correspond to the lifting of any constraints of information flow between the two areas. One caveat here is that the empirical work supporting such a model has come from simple visual attention tasks, and more work needs to be done in order to get a clearer view on the role of alpha modulation in more complex cognitive processes such as language. Nevertheless, based on this understanding, our findings are indicative of a top-down downregulation of alpha oscillations that operate as part of a broader attentional control network throughout the brain’s sensory channels, which during speech comprehension act to promote the spread of information between language relevant cortices (Sadaghiani and Kleinschmidt 2016; Alavash et al. 2021). This interpretation further fits with the wider understanding of the role of alpha oscillations in controlling the access, transfer and storage of information within the brain (Klimesch 2012; Bonnefond et al. 2017).

Mechanistically, we tentatively suggest that there is a link between the earlier decreased alpha power (observed around the onset of the second word) and the later decreased alpha phase-locking, with each reflecting a complimentary but distinct part of the neural syntax composition process. However, in order to confirm and strengthen this interpretation, future research with a design utilising more trials (i.e., to assess phase-locking in low alpha power vs. high alpha power trials) and more participants are required. With regard to the latter point, to explore correlations at the participant level between these two observed effects, a considerably large sample size would be required (recommended at least 80 participants for a stable correlation estimate in neuroimaging research; Grady et al., 2021). Addressing this question would further enhance the understanding of syntax composition in the brain.

### Summary

In sum, when comparing minimal pseudo-verb sentences for which it was highly likely that syntactic binding may plausibly occur (“*she grushes*”) to wordlists for which binding was highly unlikely to occur (“*cugged grushes*”), we first found evidence of less alpha power in left-lateralized brain regions; we suggest that this reflects an expectation of binding to occur. Second, we found that during syntactic binding there was decreased alpha phase-locking between the LIFG and the left middle/inferior temporal cortex; we suggest that this results from increased information transfer and the strengthening of the neural network involved in syntax composition. The observed oscillatory modulations (including decreased power and inter-regional coupling) occurred in brain regions in the left hemisphere known to be relevant for language processing. These regions are not uniquely selective for syntactic processing; rather, we suggest that each region plays a contributing role in syntax composition mechanisms, and that coordination between these regions (particularly the LIFG and left middle/inferior temporal cortex) is critical for successful syntactic binding. Together, our findings contribute to the wider understanding of the rapid spatial-temporal dynamics of syntactic processing in the brain that occur independently of semantic processing; we suggest that future research builds on this by exploring the mechanistic links between different oscillatory characteristics of the syntactic binding process.

## Acknowledgments

This research was supported by an Economic and Social Research Council (ESRC) studentship awarded to Sophie M. Hardy from the University of Birmingham Doctoral Training Centre (grant number: ES/J50001X/1). We thank Jonathon Winter and Nina Salman for their help with data collection. The authors report no potential conflict of interest.

## SUPPLEMENTARY MATERIALS

### SUPPLEMENTARY MATERIALS 1

**Supplementary Figure 1.**
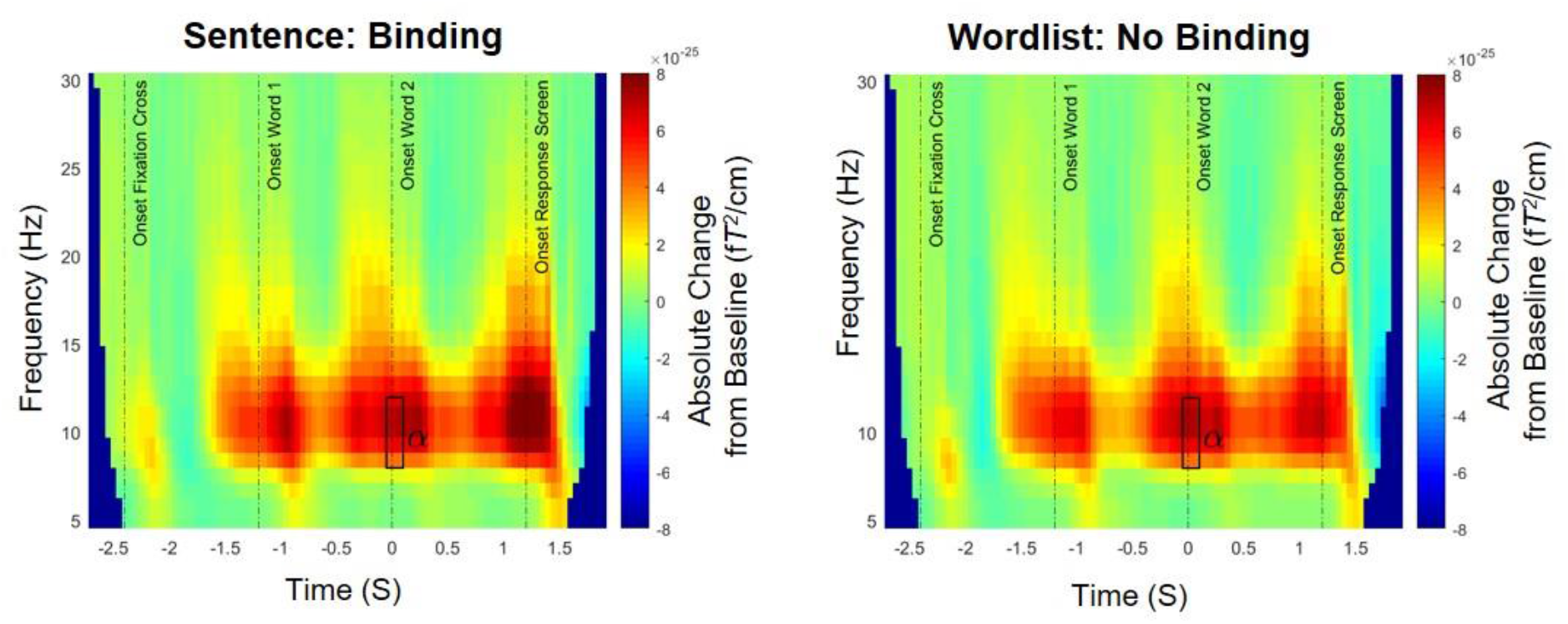
Time-frequency representations (TFRs) of power averaged across all sensors for the Sentence condition [left panel], in which syntactic binding occurred (e.g., “*she grushes*”) and the Wordlist condition [right panel], in which no binding occurred (e.g., “*cugged grushes*”). Power is expressed as an absolute change from the baseline period (−2.1s to −1.6s), which occurred during the fixation cross presentation. The complete epoch is shown. Time relates to the main time period of interest, epoched around the onset of the second word (presented at 0s). We analysed both our time period of interest (−0.5s to 1s of the epoch) and the wider timeframe of the auditory stimuli presentation (−1.3s to 1.2s). The rectangle highlights the time period showing a significant difference in alpha power (8-12 Hz) between the two conditions (−0.05s to −0.1s; *p* < .05). We did not observe any other significant effects in our time-frequency analyses.

### SUPPLEMENTARY MATERIALS 2

**Supplementary Figure 2.**
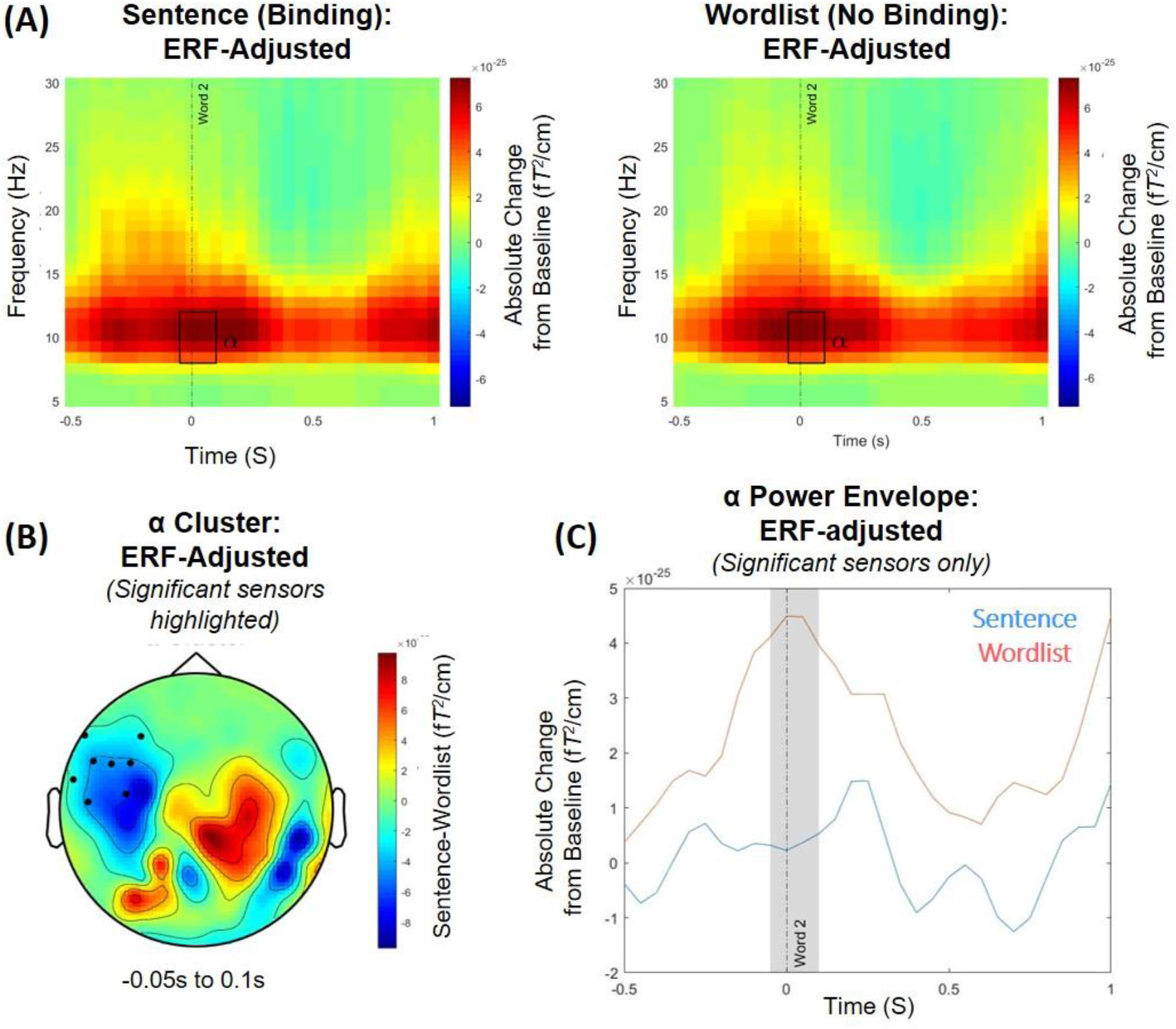
To ensure that the observed oscillatory changes were not just the spectral representation of the event-related fields (ERFs), the ERF components were subtracted from the time-frequency representation of the oscillatory data (Mazaheri & Picton, 2005). The subtraction was achieved by first generating the time frequency decomposition of the ERF data for each condition and participant separately. Next, the time frequency power spectra of the ERF were subtracted from the time frequency power spectra of the MEG signal for each condition. The subsequent power changes in the time-frequency domain were used to generate time frequency power spectra differences between experimental conditions (Sentence vs. Wordlist). We then reanalysed the ERF-adjusted TRFs of power using the same statistical methods outlined in the primary time-frequency analyses. In these ERF-adjusted analyses, we found the same significant alpha power cluster at −0.05s to 0.1 (*p* = .022). This demonstrates that our observed oscillatory effect of alpha was distinct from the evoked fields. Shown in this figure is: **(A)** TFRs of the ERF-adjusted power averaged across all sensors for the sentence condition [left panel] in which syntactic binding was highly to occur (e.g., “*she grushes*”), and the wordlist condition [right panel] in which binding was highly unlikely (e.g., “*cugged grushes*”); **(B)** the scalp topography of the condition contrast (Sentence-Wordlist) of the averaged alpha power activity in the time window showing a significant difference in power between conditions; and **(C)** the time course of the alpha power envelope for the sensors showing a significant difference in power between the Sentence (blue) and Wordlist (red) conditions.

## Footnotes

1 Following the neuroimaging binding literature, we did *not* include a behavioural judgement task within our MEG study because it is very important that there is no difference in judgement between the conditions for which the EEG/MEG signal is compared, otherwise judgement (yes/no) and condition (binding/no binding) would be conflated, making it impossible to directly compare the signatures of the two conditions. It is therefore better to remove the judgement element and just focus on the direct comparison of the MEG signature of syntactic binding vs. no binding.

2 Pseudo-verbs (root form, un-inflected): brop, crog, cug, dotch, grush, plag, plam, pob, prap, prass, satch, scash, scur, slub, spuff, stoff, trab, traff, tunch, vask.

3 To preview the time-frequency findings, we found significant effects in the alpha (8-12 Hz) frequency range in the time period surrounding the onset of the second word (−0.05s to 0.1s). We therefore analysed the alpha band in our source localization analyses. However, we did expand the time window slightly to −0.15s to 0.15s for the localization analyses as, for the DICS approach, the time window needs to be long enough to allow for sufficient frequency activity within the alpha range (recommended at least three cycles; 3/10 Hz = 0.3s).

4 For TFRs of the complete epoch (−2.7s to 1.9s), see **Supplementary Materials 1.**

